# Single-virus content mixing assay reveals cholesterol-enhanced influenza membrane fusion efficiency

**DOI:** 10.1101/2021.04.26.441491

**Authors:** Katherine N. Liu, Steven G. Boxer

## Abstract

In order to infect a cell, enveloped viruses must first undergo membrane fusion, which proceeds through a hemifusion intermediate, followed by the formation of a fusion pore through which the viral genome is transferred to a target cell. Single-virus fusion studies to elucidate the dynamics of content mixing typically require extensive fluorescent labeling of viral contents. The labeling process must be optimized depending on the virus identity and strain and can potentially be perturbative to viral fusion behavior. Here, we introduce a single-virus assay where content-labeled vesicles are bound to unlabeled influenza A virus (IAV) to eliminate the problematic step of content-labeling virions. We use fluorescence microscopy to observe individual, pH-triggered content mixing and content loss events between IAV and target vesicles of varying cholesterol compositions. We show that target membrane cholesterol increases the efficiency of IAV content mixing and decreases the fraction of content mixing events that result in content loss. These results are consistent with previous findings that cholesterol stabilizes pore formation in IAV entry and limits leakage following pore formation. We also show that content loss due to hemagglutinin fusion peptide engagement with the target membrane is independent of composition. This approach is a promising strategy for studying the single-virus content mixing kinetics of other enveloped viruses.

**Statement of Significance:** To replicate, enveloped viruses, like influenza A virus, must successfully deliver their contents to a host cell through viral membrane fusion. Most single-virus fusion assays require extensive fluorescent labeling of virions which can be perturbative to fusion kinetics. Here, we utilize content-labeled vesicles in a single-virus content mixing assay, which eliminates the need to fluorescently label virus contents. We use this assay to show that target membrane cholesterol increases the fraction of stable influenza virus content mixing events. This assay also enables the study of target membrane destabilization due to viral fusion peptide engagement.

## Introduction

The transfer of genetic information from a virus to a host cell is a critical precursor for viral replication. For enveloped viruses, in order for this transfer to occur, the viral membrane must fuse to a host membrane, which is followed by the formation of a fusion pore through which the viral genome is transferred into the target cell (1, 2). For viruses that have segmented genomes, like influenza A virus (IAV), effective genome transfer is especially vital to produce replicated virions that are fully functional.

IAV entry occurs in the endosome and is mediated by its envelope protein hemagglutinin (HA). Fusion is triggered by low pH as the endosome matures, and HA undergoes a large conformational rearrangement to insert its hydrophobic fusion peptide into the host membrane (3, 4). Kinetic studies and simulations have shown that for IAV fusion to occur, several neighboring HA fusion peptides must successfully engage with the target membrane (5–7). Once these minimal requirements have been satisfied, the membranes of influenza and host membrane mix to form a structure that is referred to as a hemifusion stalk and subsequently a hemifusion diaphragm, then a fusion pore is formed and widened to allow for genome transfer (8). While IAV and HA-mediated fusion have been widely studied, there are still questions remaining about the detailed mechanism of pore formation, specifically the timescale and dynamics of genome transfer and how the composition of the target membrane affects efficient transfer.

The kinetics of content mixing have been studied in many *in vitro*, reductionist systems that utilize model membranes as a proxy for cells. These simplified membranes enable researchers to control target membrane compositions to systematically study the impact of various membrane components on viral fusion. Cholesterol is one membrane component that is of significant interest as it is hypothesized that the cholesterol-rich regions of host membranes facilitate viral fusion peptide engagement, which can promote membrane fusion (9–11), and vesicle fusion studies have found that cholesterol enhances fusion and pore formation (12–14). In bulk studies, where fusion between virions and vesicles is observed by fluorescence dequenching and an overall increase in fluorescence signal, cholesterol speeds the rates of IAV lipid mixing (hemifusion) and content mixing (pore formation) (15). However, ensemble measurements report average behavior for a large number of particles, and the readout for content mixing can be confounded by issues like vesicle rupture, content leakage, and viral or target vesicle aggregation.

To address the shortcomings of bulk measurements, efforts have been made towards developing single-virus assays to monitor the kinetics of lipid and content mixing (6, 16–18). These single-virus assays can characterize the heterogeneity in viral fusion events and short-lived kinetic intermediates that are typically lost in bulk averaging studies. In fluorescence-based experimental architectures, virions are lipid-and/or content-labeled with self-quenched concentrations of fluorescent dyes. Content-labeling virions with sufficient amounts of dye to be useful involves a long incubation period (anywhere from 16 hrs to 2 days) in a water-soluble dye (6, 19, 20). The labeling process is specific to each virus, and it requires significant optimization to successfully incorporate dyes while ensuring that labels do not impact viral infectivity. In most prior work, labeled virions were fused to a supported lipid bilayer (SLB) formed on a solid support, where content mixing is detected as dequenching, followed by a sharp drop in fluorescence intensity as the content dye diffuses away.

While SLBs may be a suitable target for studies of outer leaflet mixing, it is not clear where a content dye goes if a pore forms and how the solid support affects engagement of the inner leaflet. Similar issues arise in models for SNARE mediated membrane fusion where tethered bilayers, held some distance from the solid support, have been used to surmount this limitation (21). Additionally, as viruses fuse to a target SLB and the membranes mix, the composition of the target membrane changes over time. This change in SLB composition makes it difficult to objectively study how target membrane composition affects viral fusion kinetics.

To address these concerns about SLB architectures, we and others have developed single-virus lipid mixing assays where viruses fuse to tethered vesicles instead of SLBs (22–24). In these assays, vesicles are tethered to labeled virions through either sialic acid receptors or synthetic DNA-lipid tethers, and single lipid mixing events are detected by fluorescence microscopy. Because individual virions are tethered to different vesicles, the target membrane composition does not change before a lipid mixing event occurs, and dye labels stay confined to each virus-vesicle pair after mixing. For IAV, it has been shown using target vesicles from 30-100 nm in diameter that the overall starting curvature of the target membrane does not affect the rate of single-virus lipid mixing (25). As long as surface passivation is optimized to prevent nonspecific binding, vesicles offer an attractive alternative to SLBs as host membrane mimics.

To avoid the process of adding a self-quenched concentration of fluorescently-tagged lipids to the viral envelope, we previously created a new version of the single-virus lipid mixing assay where the experimental readout for viral fusion is fluorescence dequenching of lipid-labeled target vesicles. Using this assay, we found that that target membrane cholesterol enhances the efficiency of influenza A virus (IAV) lipid mixing but has no effect on the rate (26). Here, we present a single-virus content mixing assay that utilizes content-labeled vesicles to eliminate the difficult process of content-labeling virions. Employing vesicles as target membranes enables the characterization of content leakage dynamics that cannot be observed in typical SLB architectures.

## Materials and Methods

### Materials

Palmitoyl oleoyl phosphatidylcholine (POPC), dioleoyl phosphatidylethanolamine (DOPE), and cholesterol (CH) were purchased from Avanti Polar Lipids (Alabaster, AL). Texas Red-1,2-dihexadecanoyl-*sn*-glycero-3-phosphoethanolamine (TR-DHPE), Oregon Green-1,2-dihexadecanoyl-*sn*-glycero-3-phosphoethanolamine (OG-DHPE), fatty acid depleted bovine serum albumin (BSA), and NeutrAvidin were purchased from Thermo Fisher Scientific (Waltham, MA). Sulforhodamine B (SRB), Sepharose CL-4B, and disialoganglioside GD1a from bovine brain (Cer-Glc-Gal(NeuAc)-GalNAc-Gal-NeuAc) were purchased from Sigma-Aldrich (St. Louis, MO). Chloroform, methanol, N-(2-hydroxyethyl)piperazine-N’-(2-ethanesulfonic acid) (HEPES) buffer, and buffer salts were obtained from Fisher Scientific (Pittsburgh, PA) and Sigma-Aldrich. Polydimethylsiloxane (PDMS) was obtained from Ellsworth Adhesives (Hayward, CA). Poly(L-lysine)-graft-poly(ethylene glycol) (PLL-g-PEG) and Poly(L-lysine)-graft-poly(ethylene glycol) biotin (PLL-g-PEG biotin) were purchased from SuSoS AG (Dübendorf, Switzerland).

### Buffers

The following buffers were used. Vesicle buffer: 10 mM NaH2PO4, 90 mM sodium citrate, 150 mM NaCl, pH 7.4. Fusion buffer: 10 mM NaH2PO4, 90 mM sodium citrate, 150 mM NaCl, Ph 5.1. HB buffer: 20 mM HEPES, 150 mM NaCl, pH 7.2. Content buffer: 30 mM sulforhodamine B (SRB), 10 mM NaH2PO4, 90 mM sodium citrate, 120 mM NaCl, pH 7.4.

### Microscopy

Epifluorescence micrographs were acquired with a Nikon Ti-U microscope using a 100X oil immersion objective, NA = 1.49 (Nikon Instruments, Melville, NY), a Spectra-X LED Light Engine (Lumencor, Beaverton, OR) for illumination and an Andor iXon 897 EMCCD camera (Andor Technologies, Belfast, UK) with 16-bit image settings. Images were captured with Metamorph software (Molecular Devices, Sunnyvale, CA). See SI Materials for additional microscopy information.

### DNA-lipid and biotin-DNA preparation

DNA-lipids (see Table S1 for sequences) used to surface-tether viruses were synthesized as previously described (27). Biotinylated-DNA was synthesized by the PAN facility at Stanford University and diluted to the desired concentration in DI water. All DNA oligos were stored at −20°C.

### Influenza virus preparation

Influenza A virus (strain X-31, A/Aichi/68, H3N2) was purchased from Charles River Laboratories (Wilmington, MA). Virus was pelleted in HB buffer by centrifugation at 21130 rcf for 50 min and resuspended in fresh HB buffer. DNA-lipids were incorporated into the IAV envelope by incubating virus sample at 4°C on ice overnight as previously described (22, 26). IAV is a BSL-2 agent and was handled following an approved biosafety protocol at Stanford University.

### Vesicle preparation

Lipid mixtures were prepared in chloroform, dried down to a film under argon gas, and the film was dried under house vacuum for at least 3 hours. Mixtures contained 10-40 mol% CH, 20 mol% DOPE, 2 mol% GD1a, and remaining mol% POPC (see Table S1). Dried lipid films were resuspended in content buffer by vortexing, and large unilamellar vesicles (LUVs) with a nominal diameter of 100 nm were prepared by extrusion. Vesicle suspensions were stored at 4 °C and used within a week. To incorporate DNA-lipids into the outer leaflet of vesicles composed of 10% CH, DNA-lipids were added to a vesicle suspension and incubated overnight at 4 °C, as described in previous studies (22, 28). Immediately before use in a content mixing experiment, vesicles were purified from free SRB dye on a CL-4B size exclusion column and equilibrated with vesicle buffer. After equilibration, vesicles were used within four hours.

### Surface and architecture preparation

The single virus content mixing architecture was prepared as described in Fig. 1. In a microfluidic flow cell (see SI Methods for details), glass slides were functionalized as previously described (26). Next, 5 µL of IAV in HB buffer (roughly 5.4 nM) displaying DNA sequence A’ (antisense to A) were introduced to the flow cell and tethered to the substrate. For experiments to vesicles with 10% CH and antibody content mixing experiments, IAV also displayed DNA sequence B (Fig. S1). After rinsing the flow cell with vesicle buffer, the surface was further passivated by incubating 10 µL of bovine serum albumin (BSA, 1 g/L) for at least 10 min to prevent non-specific binding of added target vesicles. Finally, 2-3 µL of ~100 nm diameter vesicles displaying GD1a and containing content buffer were introduced (2.8 µM nominal total lipid concentration). Vesicles with 10% CH displayed both GD1a and DNA sequence B’ (antisense to B). Content vesicles were allowed to bind for 5-10 min to control surface density and ensure spatial separation between particles. Excess unbound vesicles and BSA were removed by rinsing with vesicle buffer.

**Figure 1.**
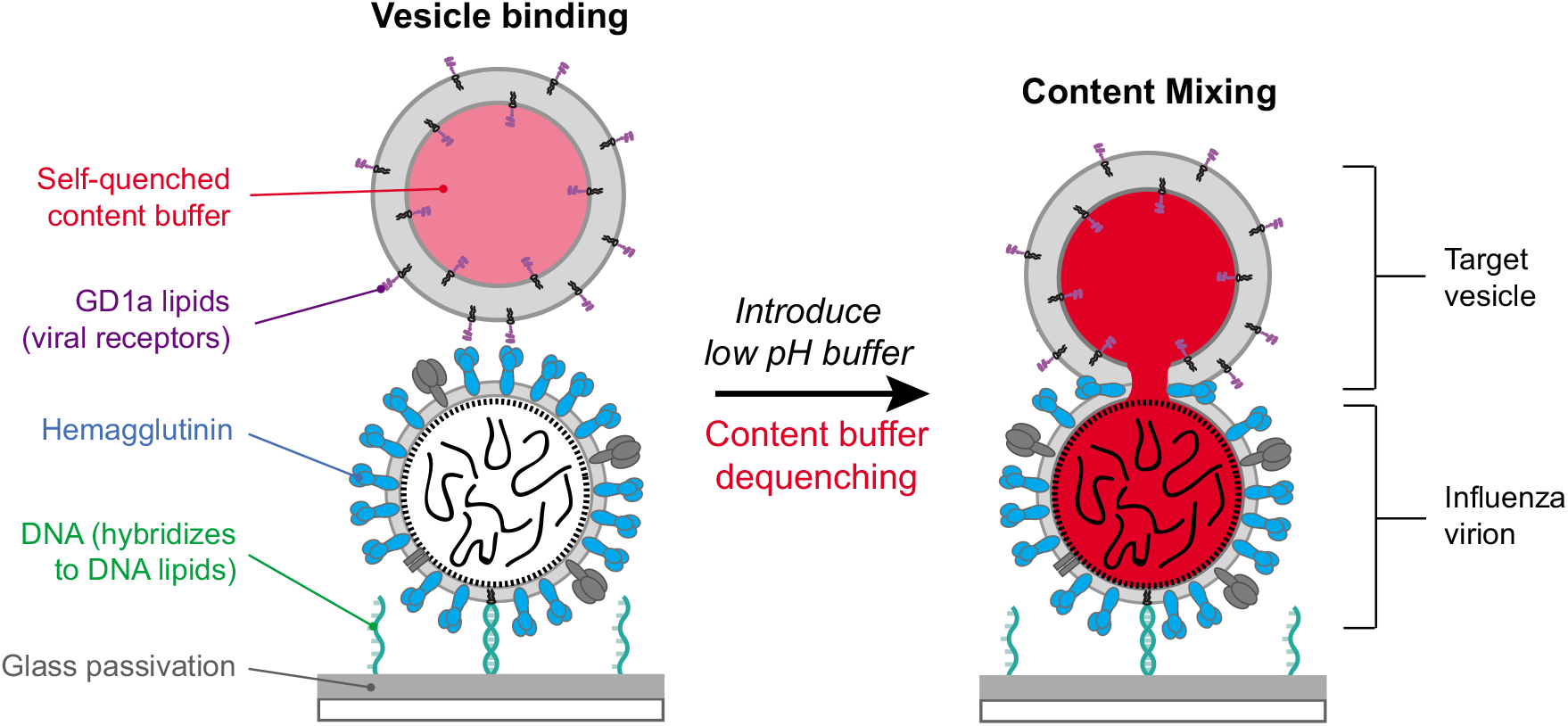
Schematic of single virus content mixing assay. Influenza A virions are tethered to a passivated (gray), DNA-functionalized glass coverslip in a microfluidic device through DNA hybridization (green). Target vesicles containing a self-quenched concentration of SRB content buffer (red) and GD1a glycolipids (purple) bind to hemagglutinin (HA, blue) of influenza A virions. For vesicles composed of 10 mol% CH, vesicles also displayed a small number of DNA-lipids that were orthogonal to the surface tethering sequences (Fig. S1). Once vesicles are bound, low pH buffer is exchanged to trigger fusion, and fluorescence dequenching occurs as the contents between target vesicle and virus mix.

### Content mixing assay

Fluorescence microscopy was used to collect a video micrograph image stream of 1200 frames at a rate of 3.47 frames/s. After the image stream was started, the pH of flow cell was rapidly exchanged from 7.4 to 5.1 using fusion buffer. In a separate experiment, tethered vesicles that contained a pH indicator (2 mol% OG-DHPE) were used to calibrate the exchange time of fusion buffer (2-3 s). The times between lowering of pH to dequenching and/or content escape were extracted using custom MATLAB (MathWorks, Inc.) scripts, as described previously (22, 26). The wait times from fluorescence traces with more than one dequenching event are excluded from cumulative distribution functions (CDFs).

### Single-virus binding assay

IAV was labeled with TR-DHPE using methods previously described (22, 29). Labeled IAV was incubated with various solutions of monoclonal antibodies for 2 hours at 22° C. SLBs (67.9% POPC, 20% DOPE, 10% CH, 2% GD1a, and 0.1% OG-DHPE) were formed through vesicle fusion. Antibody-bound virions were introduced to this bilayer and allowed to bind for 60 s. Unbound virions were rinsed away using vesicle buffer, and the resulting number of virions was quantified through spot analysis of fluorescence micrographs using MATLAB scripts. See SI Methods for more details.

## Results

### Single-virus content mixing assay

We created an assay to observe single-virus content mixing events that does not require the difficult and perturbative process of labeling viral contents (Fig. 1). First, IAV particles with no fluorescent label are incubated with an aqueous suspension of DNA-lipids. After at most a few DNA-lipids have incorporated into the IAV envelope (26), virions are tethered to a passivated glass slide in a PDMS microfluidic flow cell through DNA hybridization. It is essential that the surface be completely passivated to prevent non-specific adhesion of virus particles or vesicles. Next, target vesicles containing a self-quenched concentration of the water-soluble content dye sulforhodamine B (SRB) and displaying GD1a glycolipid receptors are introduced at a dilute concentration and allowed to bind to the tethered IAVs. Vesicles are nominally 100 nm in diameter to ensure that mixing with influenza A virions would roughly double the volume of the dye and result in dequenching. We have previously shown using dynamic light scattering that varying the amount of cholesterol in the target membrane does not significantly change the size of vesicles (26). We note that recent evidence has shown that extrusion yields a fraction of vesicles that are multilamellar (30), although not under our preparation conditions and for these specific compositions.

We observed that vesicles composed of 20-40 mol% cholesterol and 2 mol% GD1a bind stably to tethered IAV. However, the interaction of vesicles composed of 10 mol% CH with IAV is reversible and transient, which causes vesicles to become unbound during the process of rapid buffer exchange and so they cannot be reliably monitored. The reversible binding of 10 mol% CH vesicles, but stable binding of 20-40 mol% CH vesicles, is consistent with previous work which showed that higher CH levels increase IAV binding avidity, likely due to CH/GD1a cluster formation (31). Therefore, to stably tether 10 mol% CH target vesicles, we added an orthogonal DNA-lipid to target vesicles to hybridize with a complementary sequence in the IAV envelope (sense/antisense sequence B, see Table S1 and Fig. S1). As previously shown, DNA-lipid incorporation into vesicles and the IAV envelope does not follow Poisson statistics (22, 26, 28). Of vesicles that display at least one DNA-lipid, the median number incorporated is 4 DNA-lipids per vesicle (Fig. S1).

Once content-labeled vesicles are bound to IAV at a high, yet still spatially resolvable density (detected by their weak residual SRB fluorescence), unbound vesicles are rinsed from the flow cell using a low flow rate to prevent bursting or leakage of content-labeled vesicles. Next, the pH of the flow cell is exchanged from pH 7.4 to 5.1, occurring over 2-3 s (calibrated in virus-free samples using OG-DHPE labeled vesicles as a pH sensor). This rapid buffer exchange is also conducted at a flow rate that does not lead to leakage or bursting of content-labeled vesicles. The fluorescence intensity, or SRB signal, of each content-labeled vesicle is monitored over time through a video micrograph collected for 1200 frames at 288 ms/frame.

Upon lowering the pH, we observed several types of fluorescence time traces and classified them into four categories: content mixing (Fig. 2A), content mixing followed by content loss (Fig. 2B), content loss (no evidence of mixing, Fig. 2C), and no change. For the majority of traces (70-80%), no change was detected. We describe the first three categories in more detail below.

**Figure 2.**
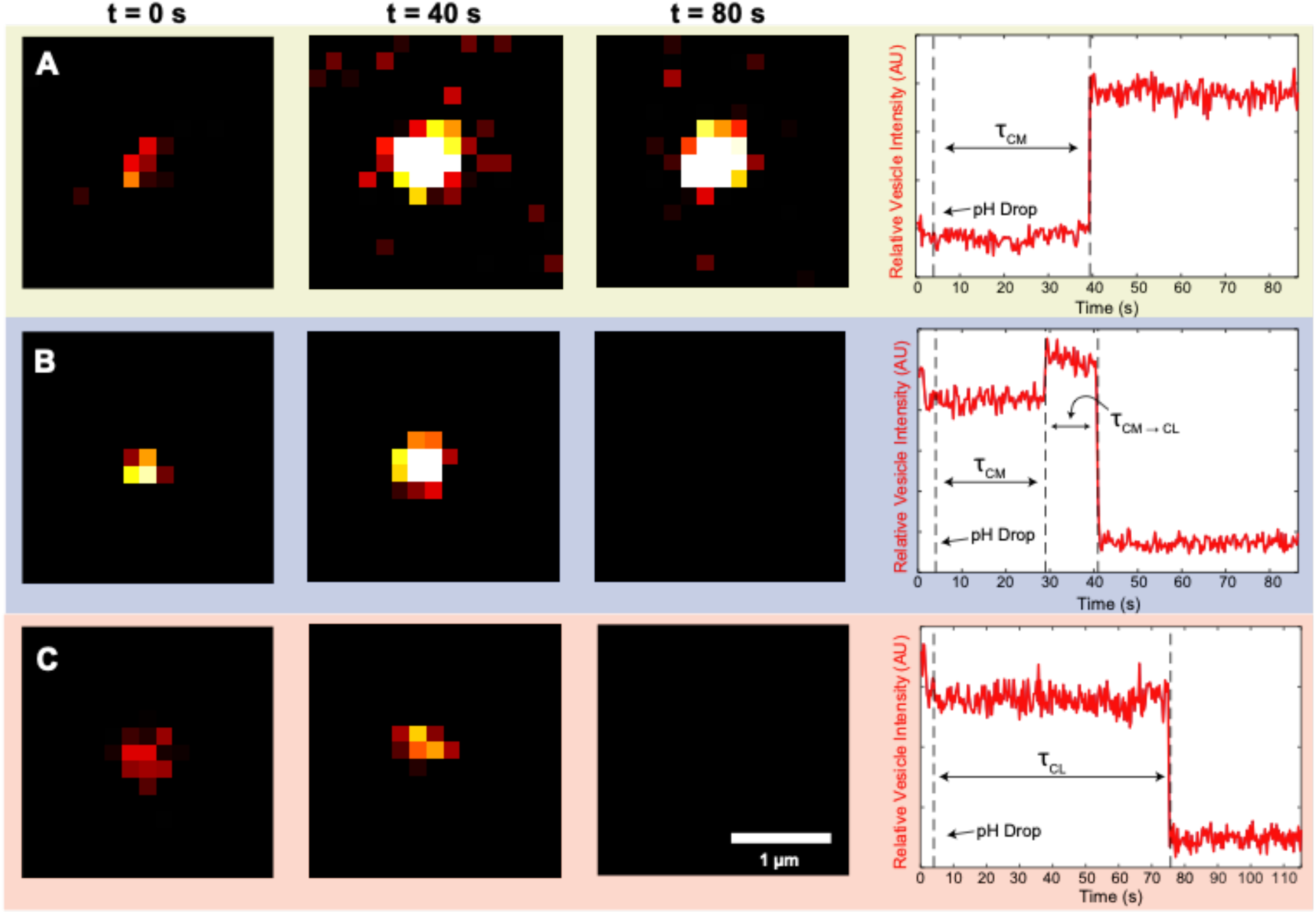
Lowering the pH triggers virion-vesicle content mixing detected by fluorescence dequenching and content loss indicated by dye disappearance. When the pH is lowered, some vesicles exhibit a change in fluorescence intensity; these traces were classified into three categories. Example microscope images of individual content-labeled vesicles bound to surface-tethered influenza virions (left). At pH 7.4, content-labeled vesicles are dim but detectable. Representative observed fluorescence intensity time traces correspond to single content-labeled vesicles (right). A) Content mixing: after the pH is lowered to 5.1, the vesicle displays dequenching due to dilution of contents with the interior of the influenza virion. Following content mixing, the fluorescence intensity of the vesicle-virion complex stays constant. B) Content mixing followed by content loss: after the pH is lowered to 5.1, the vesicle displays dequenching due to content mixing. After the dequenching event, there is a sustained period of constant fluorescence, followed by content loss, as indicated by a sharp decrease in fluorescence intensity. For both A and B, the wait time is defined as τ_CM_, or the time from the pH drop to the content mixing event. The time from content mixing to loss, or τ_CM_ → τ_CL_, is defined as the time from the content mixing event to content loss. C) Content loss, no detectable mixing: after the pH is lowered to 5.1, no content mixing event is detected, and the content-labeled vesicle undergoes content loss. For C, content dequenching occurs more quickly than the collection frame rate (288 ms/frame). The time to content loss, or τ_CL_ is defined as the time from pH drop to content loss.

In a content mixing trace, after the pH is lowered to 5.1, the vesicle displays dequenching due to dilution of contents with the interior of the influenza virion. This interpretation of content mixing is consistent with previous studies of vesicle fusion between two vesicles (32). After the content mixing event, the fluorescence intensity of the vesicle stays constant (Fig. 2A). Roughly 3-15% of traces were classified as content mixing, which is described in more detail below.

We describe the second category as content mixing followed by content loss, which made up 1.0-1.5% of traces. After the pH is lowered to 5.1, these vesicles display dequenching due to content mixing. After the dequenching event, there is a sustained period of constant fluorescence, followed by content loss, as indicated by a sharp decrease in fluorescence intensity (Fig. 2B). For all vesicles that undergo content mixing and content mixing followed by content loss, the wait time is defined as τ_CM_, or the time from the pH drop to the content mixing event. The interval after mixing where the fluorescence intensity stays constant is defined as τ_CM→CL_, or the time from the content mixing event to content loss.

We refer to the final category of time traces as content loss only (no mixing). In these vesicles, after the pH is lowered to 5.1, no increase in fluorescence characteristic of content mixing dequenching is detected, and there is a sharp decrease in fluorescence intensity, which corresponds to content loss (Fig. 2C). The time to content loss is defined as τ_CL_, or the time from pH drop to content loss. For most content loss traces, content dequenching occurs on a timescale that is faster than the video micrograph collection frame rate (288 ms/frame).

Content mixing, with and without subsequent content loss, is only observed at low pH in the presence of IAV. At neutral pH, we did not observe any instances of content mixing followed by content loss (Fig. S2). When content-labeled vesicles are tethered to glass through DNA-hybridization in the absence of IAV, we do not observe any content mixing, or content mixing followed by content loss (Fig. S2).

### Cholesterol enhances the efficiency of content mixing but not the rate

To understand how target membrane cholesterol affects the rate of content mixing, we took τ_CM_ values for each target vesicle composition and plotted them in individual cumulative distribution functions (CDFs, Fig. S3). CDFs contain τ_CM_ values from vesicles that underwent content mixing, as well as content mixing events that were followed by content loss. When we compared the CDFs for each composition, the rates of content mixing do not change significantly as target membrane cholesterol increases (Fig. 3A).

**Figure 3.**
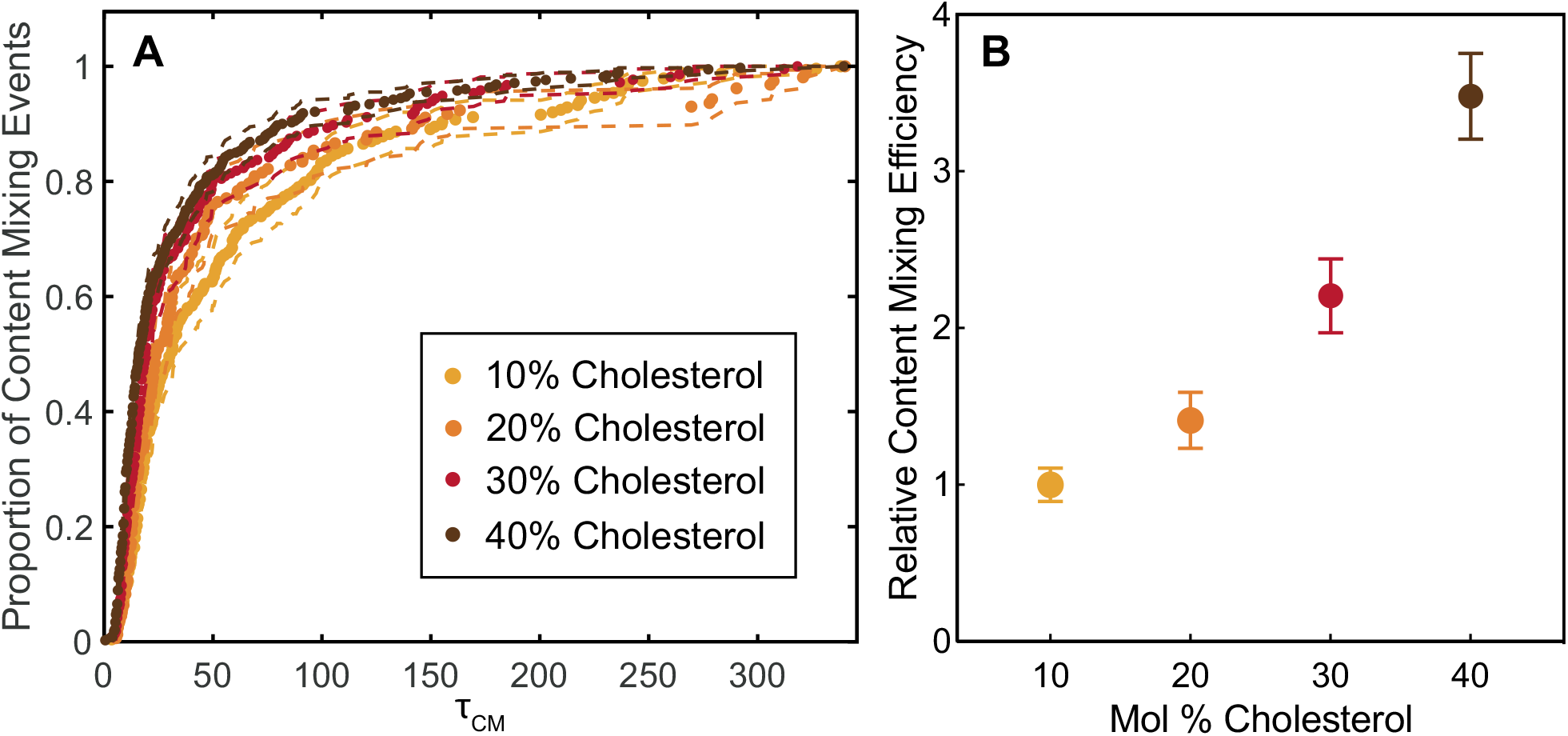
Cholesterol enhances the efficiency of single-particle IAV content mixing but does not affect the rate. A) Wait times from pH drop to content mixing (τ_CM_) for individual content mixing events are plotted as cumulative distribution functions (CDFs). From 10 to 40 mol% cholesterol, the rate of IAV content mixing is the same within bootstrap resampling error (95% confidence intervals, dashed lines). B) The relative efficiency, or fraction of target vesicles that undergo content mixing, increases 3.5-fold as the mole percentage of cholesterol in target vesicles increases. The content mixing efficiency to membranes with 10 mol% cholesterol is normalized to 1, and points represent the average efficiency value ± bootstrap resampling error. The number of content mixing events for 10, 20, 30, and 40 mol% cholesterol is 238/5250, 157/2458, 215/2153, and 380/2411, respectively. The total number of vesicles analyzed is proportional to the number of individual experiments executed, and not due to inherent differences in binding behavior. Kinetic data for each composition were compiled from at least four different independent viral preparations.

The number of vesicles that undergo content mixing, or NCM, can be divided by the total vesicles in a field of view (FOV), or N, to yield the content mixing efficiency. Target membrane cholesterol enhances the efficiency of content mixing (Fig. 3B). For all target membrane compositions tested, content mixing was a slower process than lipid mixing (see Fig. S4 for direct comparisons of content mixing and previously published lipid mixing rates).

### Cholesterol increases the number of content mixing events that result in stable pore formation

We quantified the rate and frequency of content mixing events that were followed by content loss. The frequency was calculated by dividing the number of content mixing traces that result in content loss, or N_CM→CL_, by the total number of vesicles that undergo content mixing, or N_CM_. We compared this fraction, N_CM→CL_/N_CM_, for vesicles composed of 10-40 mol% cholesterol and found that target membrane cholesterol decreases the fraction of content mixing events that result in eventual content loss (Fig. 4A). The τ_CM→CL_ intervals for each composition were plotted in individual CDFs and compared, but N_CM→CL_ values for high cholesterol containing vesicles were too low to observe any significant difference between compositions (Fig. S5).

**Figure 4.**
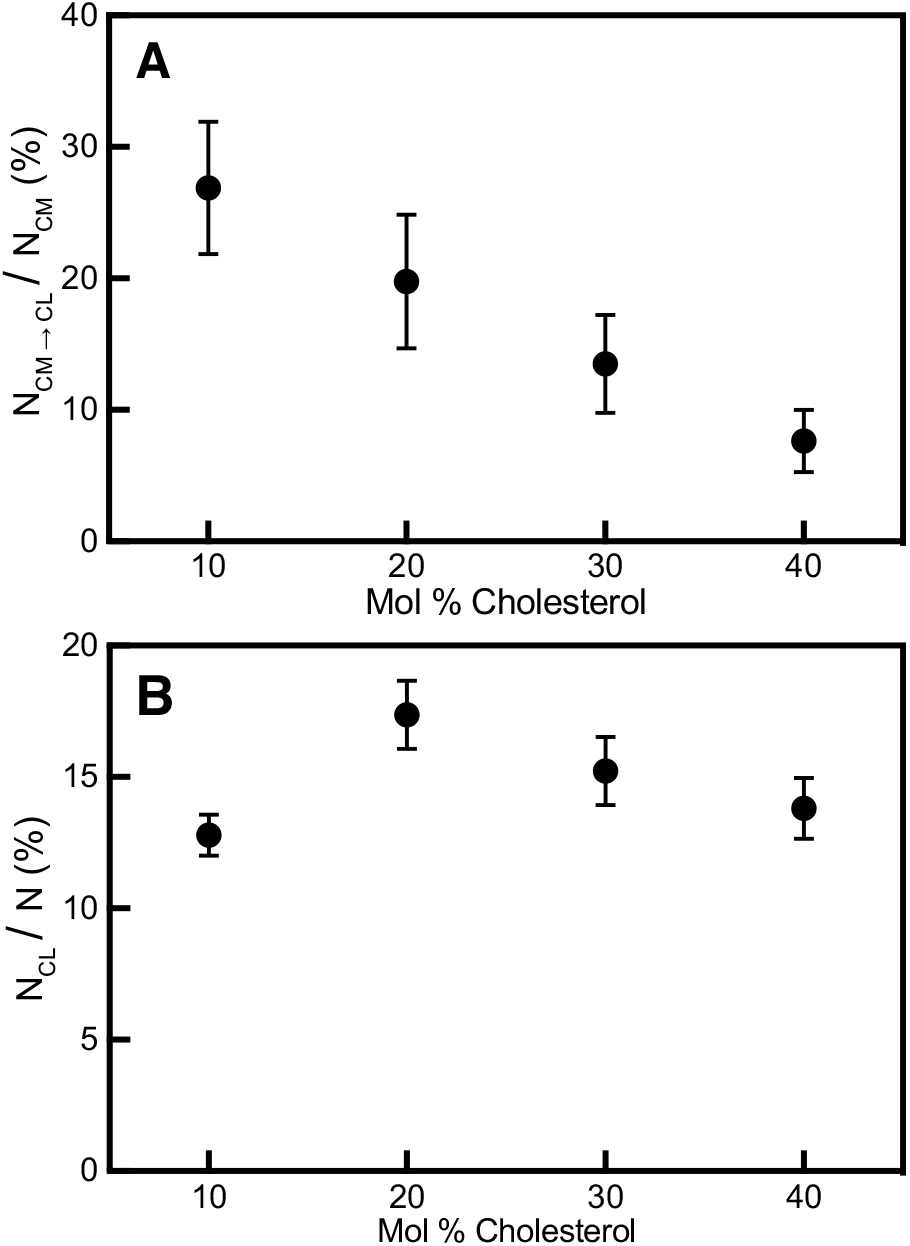
Cholesterol decreases the number of content mixing events that result in content loss but has no effect on the frequency of content loss. A) Target membrane cholesterol decreases the fraction of content mixing events that result in content loss. The fraction of content mixing events that result in content loss for 10, 20, 30, and 40 mol% cholesterol is 64/238, 31/157, 29/215, and 29/380, respectively. B) Increasing target membrane cholesterol has no significant effect on the fraction of vesicles that undergo content loss with no previous mixing event. The number of content loss events for 10, 20, 30, and 40 mol% cholesterol is 724/5250, 427/2458, 328/2153, 333/2411, respectively. Points represent the average fraction ± bootstrap resampling error.

### Kinetics of content loss are independent of membrane composition

Next, we analyzed the rate and frequency of vesicles that displayed content loss (with no content mixing event detected) for each target membrane cholesterol composition. Content loss is a competing process to content mixing; in our assay, once vesicles undergo content loss, we are unable to monitor whether a productive mixing event occurs later. As a result, we sought to understand whether the frequency of content loss events was cholesterol dependent, which could be a confounding factor in content mixing and loss efficiencies.

The frequency of content loss events was calculated by dividing the number of vesicles that were classified as content loss, NCL, by the total number of vesicles monitored, N. When we compared the frequency of content loss, NCL/N, for vesicles composed of 10-40 mol% cholesterol, we found that altering target membrane cholesterol does not have a significant effect (Fig. 4B).

The τ_CL_ intervals for each composition were plotted in individual CDFs (Fig. S6) and compared, and cholesterol also did not have a significant effect on the rate of content loss. These findings indicate that the process that leads to content loss is not affected by membrane cholesterol composition. Although we do detect a few content loss events when the vesicles are exchanged with neutral pH buffer, there are significantly more content loss events at low pH when content-labeled vesicles are tethered to IAV (Fig. S7). Due to this background of content loss events that take place when flow cells are changed with neutral pH buffer, all experiments were rinsed with a standardized amount of buffer.

Since content loss was independent of target membrane composition, we hypothesized that the events could be derived from IAV HA engagement with target membranes. To test the role of HA engagement in content loss, we neutralized HAs by introducing monoclonal antibodies to bind to the HA receptor binding domain. Antibodies were not fluorescently labeled; the extent of antibody coverage was determined through a functional single-virus binding assay based on previous studies (24, 31) where virions were labeled with TR-DHPE and incubated with various concentrations of antibody solutions. Virions were then introduced to a microfluidic flow cell containing a supported lipid bilayer (SLB) that was composed of 2 mol% GD1a, and the number of virions bound was quantified through spot analysis of fluorescence micrographs. We determined that incubating IAV with two concentrations of monoclonal antibodies, 0.05 and 0.5 mg/mL, led to a roughly 50% and 100% reduction of IAV binding in comparison to IAV with no antibodies (Fig. S8).

Next, we utilized the single-virus content mixing assay to understand how neutralizing HAs through antibody binding affects content mixing and content loss. Because antibodies prevent IAV from binding to GD1a, target vesicles were bound to IAV through DNA-lipid hybridization (median of 4 DNA-lipids/vesicle, Fig. S1). Antibodies decreased the content mixing efficiency of IAV to target vesicles composed of 10 and 40 mol% cholesterol (Fig. S9). Additionally, neutralizing HA with antibodies decreased the frequency of content loss events for both target vesicle compositions (Fig. 5). The reduction in content mixing efficiency is consistent with previous studies that showed that neutralizing antibodies decrease the efficiency of IAV lipid mixing (42).

**Figure 5.**
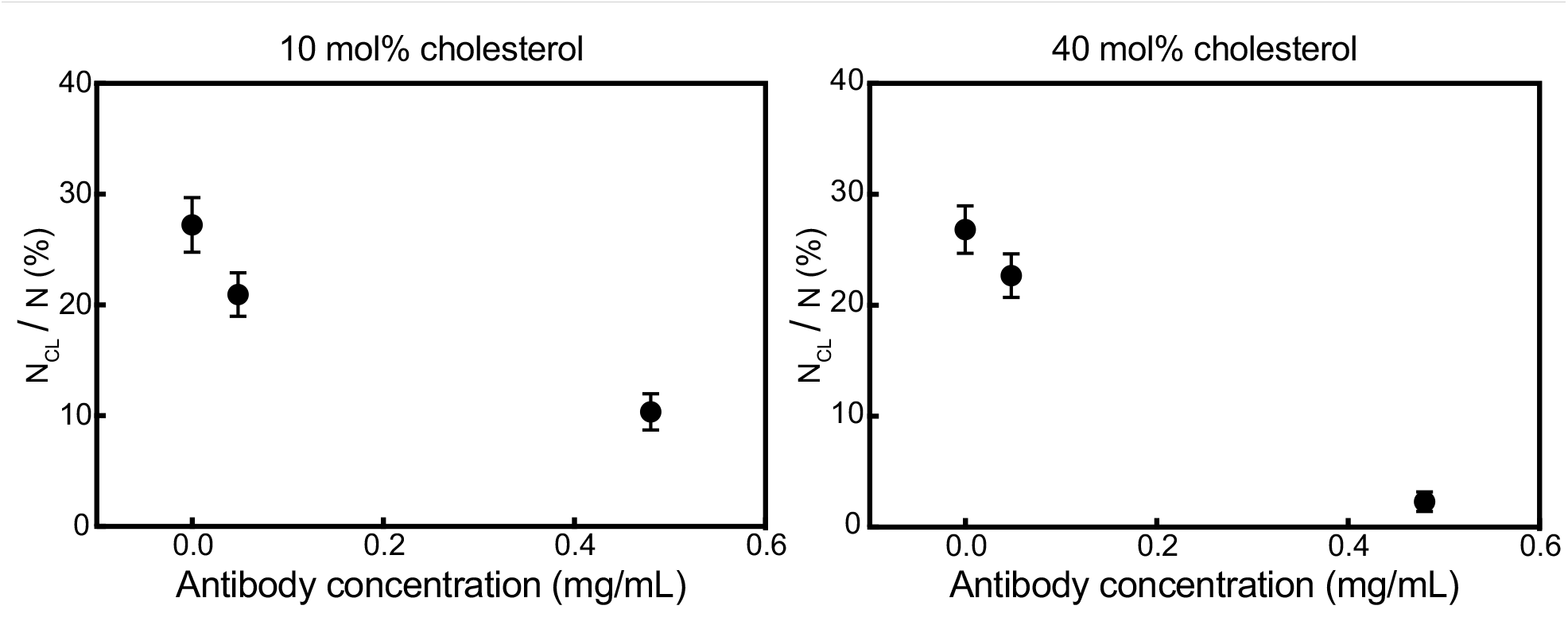
IAV antibodies decrease the frequency of content loss events to membranes containing 10 and 40 mol% cholesterol. Influenza A virions were incubated with various concentrations of monoclonal antibodies. Content-labeled vesicles with 10 and 40 mol% cholesterol were tethered to virions through DNA-lipid hybridization, and the pH was lowered to trigger content mixing and content loss. For both target membrane compositions, the addition of antibodies to IAV decreased the frequency of content loss events. Each data point represents the average frequency of content loss ± bootstrap resampling error for at least 900 vesicles.

## Discussion

We designed a single-virus content mixing assay that employs content-labeled vesicles as target membranes and does not require the process of labeling viral contents. By systematically altering the composition of vesicles used in this assay, we found that cholesterol enhances the efficiency of IAV content mixing but has no effect on the rate. The effect of target membrane cholesterol on content mixing is unsurprising given our previous finding that cholesterol also increases the efficiency of lipid mixing and has no effect on the rate (26). If there is just one rate determining step between IAV lipid mixing and content mixing as is widely believed (6), it follows that the rate of content mixing would be similarly unaffected by target membrane cholesterol. The enhancement in content mixing efficiency due to cholesterol is consistent with other IAV fusion studies that varied target membrane composition (15), and also further supports the hypothesis that the negative spontaneous curvature of cholesterol stabilizes the formation of the highly curved structure that is necessary for pore formation and widening (33, 34).

Cryo-electron tomograms of IAV fusing to vesicles have suggested that the negative spontaneous curvature (SC) of target membrane cholesterol acts as a “pathway switch,” where membranes with less than 31 mol% cholesterol go through a “rupture-insertion” pathway as opposed to the canonical, non-leaky hemifusion stalk pathway (35). A subsequent study utilized a poration assay to expand on this theory by monitoring the influx of dye into GUVs during IAV fusion. The assay concluded that the trend in membrane rupture was generalizable to target membrane SC; when SC was below −0.20 nm^−1^, fusion was predominantly non-leaky, while above that threshold, most fusion events were leaky (36). Our observations are generally in agreement with the hypothesis that the negative SC of cholesterol decreases the number of content mixing events that lead to content loss. However, our results suggest that increasing cholesterol stabilizes fusion pore formation in a more gradual manner, rather than a dramatic shift when SC is below −0.20 nm^−1^ (or above 20 mol% cholesterol, see ref. (26) for SC calculations of compositions used in this study). We hypothesize that our results diverge from previous studies due to differences in assay architecture. In the content mixing assay we present here, we monitor the real-time kinetics of content mixing and content loss of single fusion events, whereas in the GUV poration assay, multiple virions could fuse to one vesicle, and fusion events were not counted until 10 min after low pH was introduced.

Conductance measurements have suggested that membrane permeability temporarily increases in a composition-dependent manner during the process of fusion pore formation, which could lead to content loss as HA rearranges within fusing membranes (37). Additionally, computational simulations of vesicle fusion propose that cholesterol decreases the water penetrability of the membrane to inhibit leaky pore formation (38).

From a viral fitness perspective, unsuccessful pore formation is undesirable because it can inhibit viral replication. However, it has been shown that influenza genome transfer for single virions is often incomplete (39), and one contributing reason may be that fusion pores are more unstable when they are formed in host membrane regions that are relatively low in cholesterol. Other enveloped viruses have been shown to preferentially fuse to the boundary of cholesterol-rich regions of host membranes (40, 41), which suggests that protein-mediated pore formation for IAV may also selectively occur within or at the edges of cholesterol-enriched regions of the membrane. In this study, the target vesicle compositions do not lead to microscale domain formation and are visually homogenous through fluorescence microscopy, although the mixtures that we use have not been studied for any nanoscale phase separation. We are unaware of any method that can directly access the exact membrane composition at the areas where fusion pores are formed and determine how this correlates with function.

Finally, it is clear from our observations that the low-pH form of HA plays a key role in IAV-facilitated content loss from content-labeled vesicles, and this action is inhibited by monoclonal antibody binding. However, since there is no way to directly characterize the structure of each individual virion-vesicle pair whose kinetics are observed, it is difficult to make any conclusions about the structural basis for content loss. In other ensemble and structural studies, HA fusion peptide engagement has been hypothesized to lead to the deformation of host membranes (36, 43–47), and it is known that hydrophobic peptide insertion can lead to membrane disruption. This process could potentially create membrane gaps large enough for the content dye to escape from the vesicles in our assay.

## Conclusions

Here, we present a new strategy for observing single-virus content mixing that does not require the difficult, variable, and potentially damaging process of content labeling virions. By tethering content-labeled vesicles to IAV, we are able to characterize the effect of target membrane cholesterol on the kinetics of viral content mixing and content loss. We show that like IAV lipid mixing, the rate of IAV content mixing is not significantly changed by target membrane cholesterol, but cholesterol increases the efficiency of fusion pore formation. We also demonstrate that the engagement of the HA fusion peptide with target membranes can lead to content loss in a manner that is independent of membrane composition.

The content mixing assay we present here could be used for studying the dynamics of pore formation for other enveloped viruses where the viral receptor is known and can be reconstituted in a target vesicle, without requiring the process of labeling viral contents. One limitation of our study is that the content dye SRB is a much smaller molecule than the IAV genome. Directly detecting viral genome transfer in model systems is challenging but efforts to do this are ongoing.

## Author Contributions

KNL designed experiments, performed experiments, analyzed data, and co-wrote the article. SGB designed experiments and co-wrote the article.

## Acknowledgments

The authors thank Dr. Elizabeth Webster, Prof. Robert Rawle, and Prof. Peter Kasson for helpful discussions. Monoclonal antibodies were a kind gift from Dr. Robert Webster (St. Jude’s Children’s Research Hospital). KNL was supported by a National Science Foundation Graduate Research Fellowship. This work was supported by National Institutes of Health Grant R35 GM118044 to SGB.

