## Supplemental Information for "Single-virus content mixing assay reveals cholesterol-enhanced influenza membrane fusion efficiency"

K. N. Liu and S. G. Boxer

Supplementary Methods

*Flow cell and glass coverslip preparation.* Polydimethylsiloxane (PDMS) flow cells were prepared and plasma bonded to glass coverslips, as previously described (1). Briefly, glass coverslips (24 x 40 mm, No 1.5, VWR International, Radnor, PA) were cleaned by heating in 1:7 diluted 7X detergent in DI for 30 min, then rinsed extensively in DI water for 4 hours. Glass surfaces were annealed by baking slides in a kiln at 400℃ for 4 hours. PDMS flow cells (with channel dimensions 2.5 mm x 13 mm x 70 μm) and cleaned coverslips were plasma cleaned for at least 2 min and bonded together.

*Surface functionalization.* Glass slides were functionalized as previously described (2). Briefly, a 19:1 mixture of PLL-g-PEG (1 g/L) and PLL-g-PEG biotin (1 g/L) in HB buffer was incubated in flow cell for at least 30 min. Flow cells were rinsed with DI water and vesicle buffer, and stored overnight at 4˚C. The following day, neutrAvidin (1 g/L) was incubated for 15 min. After rinsing away excess neutrAvidin with vesicle buffer, biotin-DNA (178 μM, sequence A – see Table S1 for sequences) was incubated for 20 min. The flow cell was thoroughly rinsed with vesicle buffer to remove excess biotin-DNA.

*Additional microscopy information.* Sulforhodamine B images were obtained using a Texas Red filter cube (ex = 562/40 nm, bs = 593 nm, em = 624/40 nm) and additional excitation (560/55 nm) and emission (645/75 nm) filters. All images and video micrographs were captured with a frame rate of 288 ms/frame.

*Viral labeling.* IAV was labeled with TR-DHPE using methods previously described (1, 2). Briefly, TR-DHPE (0.75 g/L in ethanol) was mixed with HB buffer in a 1:40 ratio. 18 μL of IAV (2 mg/mL) was mixed with 72 μL of TR-DHPE/HB buffer solution and incubated for 2 hours at room temperature while rocking to incorporate fluorescently tagged lipids into IAV envelope. To separate unincorporated dye from labeled virions, about 1.35 mL of HB buffer was added and virus was pelleted by centrifugation at 21,130 rcf for 1 hour. The pellet that contained labeled virions was resuspended in 100 μL of fresh HB buffer.

*Supported lipid bilayer (SLB) formation.* SLBs were formed through vesicle fusion; 7 μL of vesicles (67.9% POPC, 20 % DOPE, 10% CH, 2% GD1a, and 0.1% OG-DHPE, conc. 0.56 mM) were added to a flow cell and incubated for 20-30 min to allow for SLB formation. Flow cells were subsequently rinsed at 1000 μL/min with at least 1 mL of DI water and 2 mL of vesicle buffer.

**Table S1. DNA-lipid sequences.**

| Name | DNA sequence (5'-3') | Location |
|---|---|---|
| A | Biotin-TTT TTT TTT TTT TTT TTT TTT TTT | Glass slide |
| A' | Lipid-AAA AAA AAA AAA AAA AAA AAA AAA | Influenza envelope |
| B | Lipid-CCC CCC CCC CCC CCC CCC CCC CCC CCC CCC CCC CCC CCC CCC CCC CCC CCC CCC CCC CCC CCC CCC CCC TCG ACA CGG AAA TGT TGA ATA CTA | Influenza envelope |
| B' | Lipid-TAG TAT TCA ACA TTT CCG TGT CGA | Target vesicle (only vesicles with 10 mol% CH) |
| X | Lipid-TGC GGA TAA CAA TTT CAC ACA GGA-AF546 | Vesicles (for DNA-lipid quantification) |

Supplementary Results

*Quantification of DNA-lipid incorporation in vesicles.* The DNA-lipid incorporation into vesicles (67.9% POPC, 20% DOPE, 10% cholesterol, 2% GD1a, 0.1% OG-DHPE), was quantified by using an AlexaFluor-546 (AF546) labeled DNA-lipid (Sequence X, Table S1). The number of DNA-lipids in each vesicle was measured by quantitative fluorescence imaging, which is described in detail in previous studies (1, 3). Briefly, the average fluorescence intensity of one AF546 DNA-lipid was determined by single step photobleaching. AF546-labeled DNA-lipids were incorporated into vesicles, then vesicles were non-specifically bound to a glass slide in a PDMS microfluidic flow cell. After the flow cell was thoroughly rinsed, vesicles were imaged in the AF546 and OG-DHPE channels. The number of DNA-lipids in each vesicle was calculated by dividing the overall AF546 intensity by the average fluorescence intensity of one AF546 DNA-lipid.

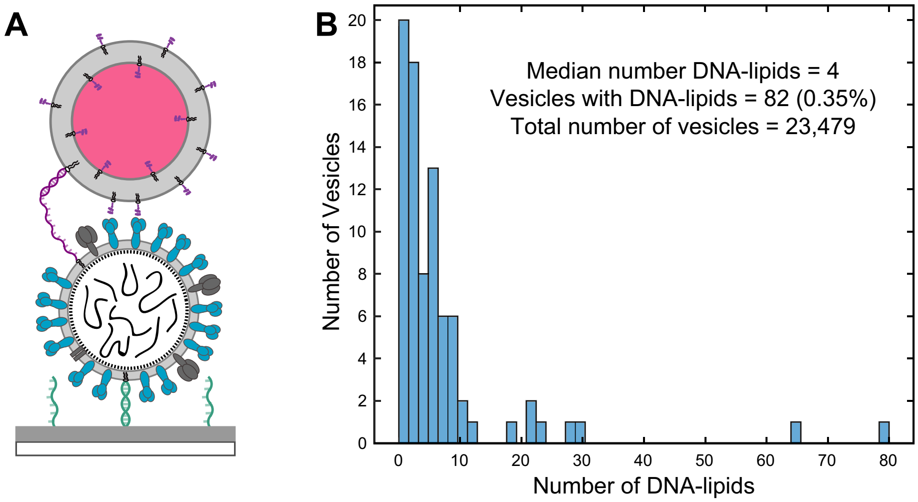


**Figure S1. Distribution of DNA-lipids in 10% CH vesicles that contained at least one DNA-lipid.**
A) Schematic of content mixing assay for vesicles containing 10 mol% CH, where in addition to GD1a, vesicles display DNA-lipids (purple, sequence B', Table S1), which hybridize to complementary DNA-lipids in IAV (sequence B). DNA-lipids were also used to tether IAV to vesicles in content mixing experiments that involved antibodies. B) Most vesicles did not display any DNA-lipids; the average number of DNA-lipids incorporated was 0.25/vesicle. Of the vesicles that displayed at least one DNA-lipid, the median number of DNA-lipids incorporated was 4. Total vesicles analyzed = 23,479.

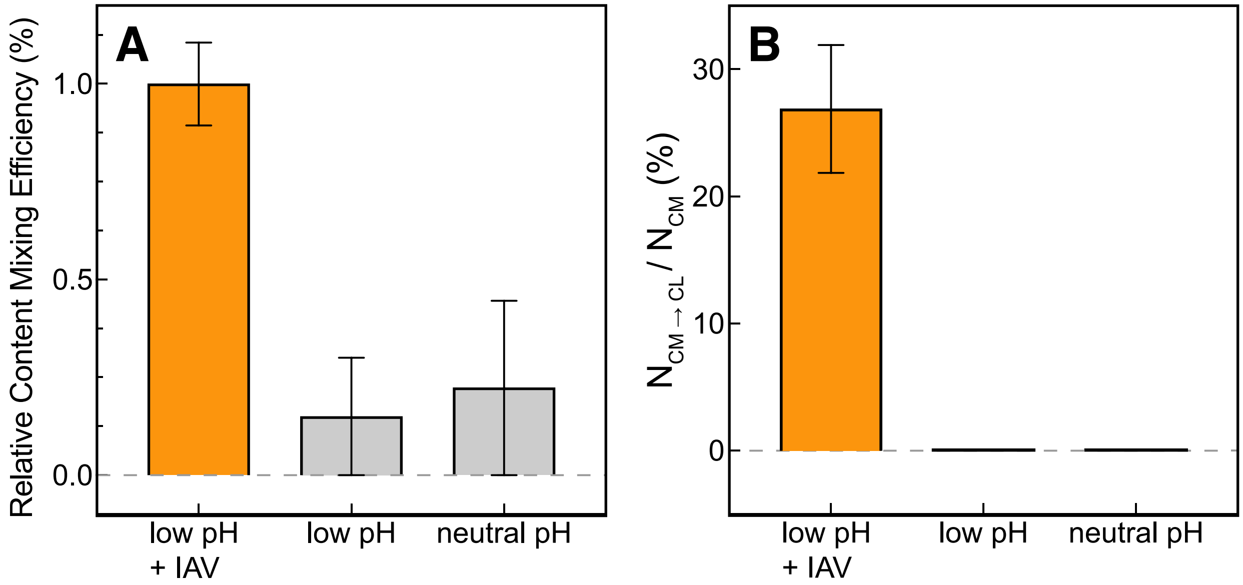


**Figure S2. Content vesicle dequenching only occurs at pH 5.1 in the presence of influenza A virus.**

In microfluidic flow cells, glass slides were functionalized as described in SI methods. For negative control experiments (gray), content-labeled vesicles (70% POPC, 20% DOPE, 10% CH) displayed DNA-lipids (sequence A') for surface tethering. Points represent the average efficiency ± bootstrap resampling error. A) In the absence of IAV, there are a few content mixing events detected, but the number falls within error of the scripts used for analysis. The content mixing efficiency at low pH for IAV-bound vesicles is normalized to 1. B) In the absence of IAV, there are no content mixing events followed by content loss.

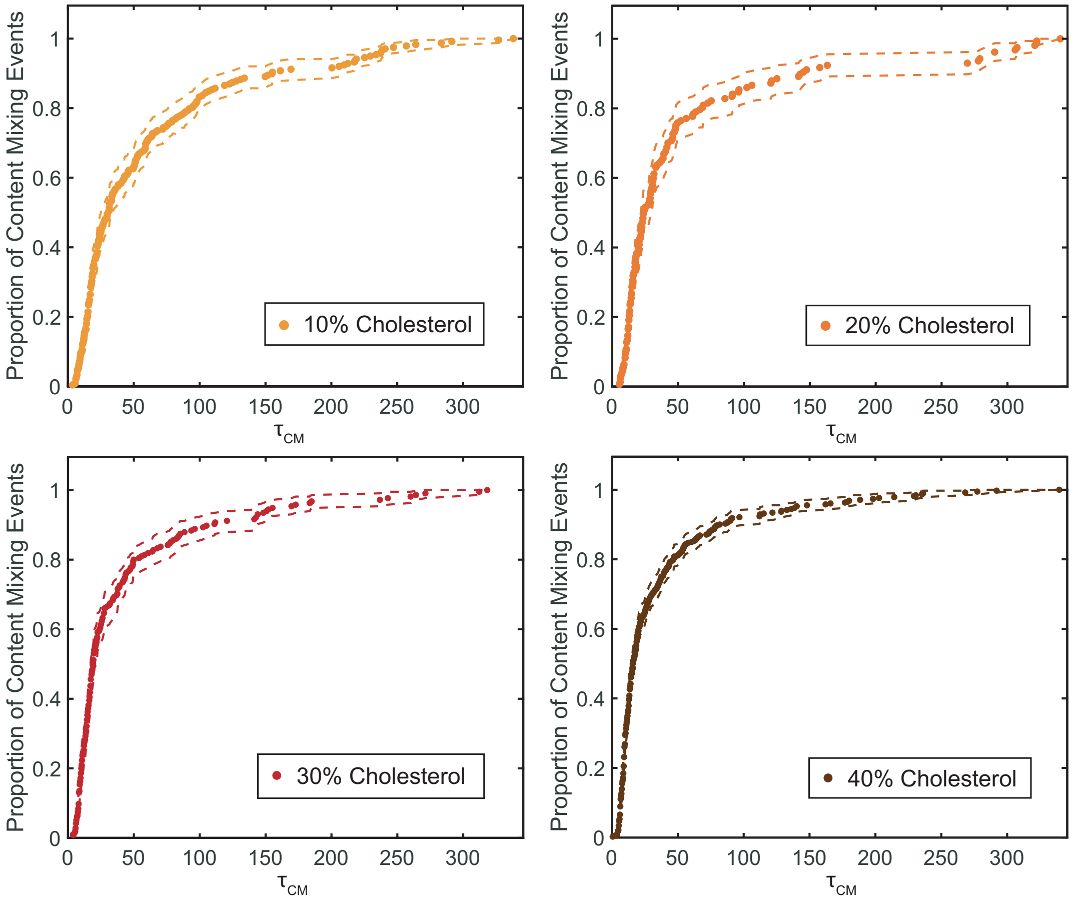


**Figure S3. Individual CDFs for IAV content mixing to target vesicles with varying cholesterol compositions (merged in Fig. 3).**

As in Fig. 3, wait times from pH drop to content mixing ($\tau_{CM}$) for individual content mixing events are plotted as CDFs. Dashed lines represent bootstrap resampling error (95% confidence intervals). As the mol fraction of cholesterol (CH) in target vesicles increases, the efficiency also increases ~3.5-fold: 238/5250 for 10% CH, 157/2458 for 20% CH, 215/2153 for 30% CH, and 380/2411 for 40% CH. The total number of vesicles analyzed is proportional to the number of individual experiments executed, and not due to inherent differences in binding behavior. Kinetic data for each composition were compiled from at least four different independent viral preparations.

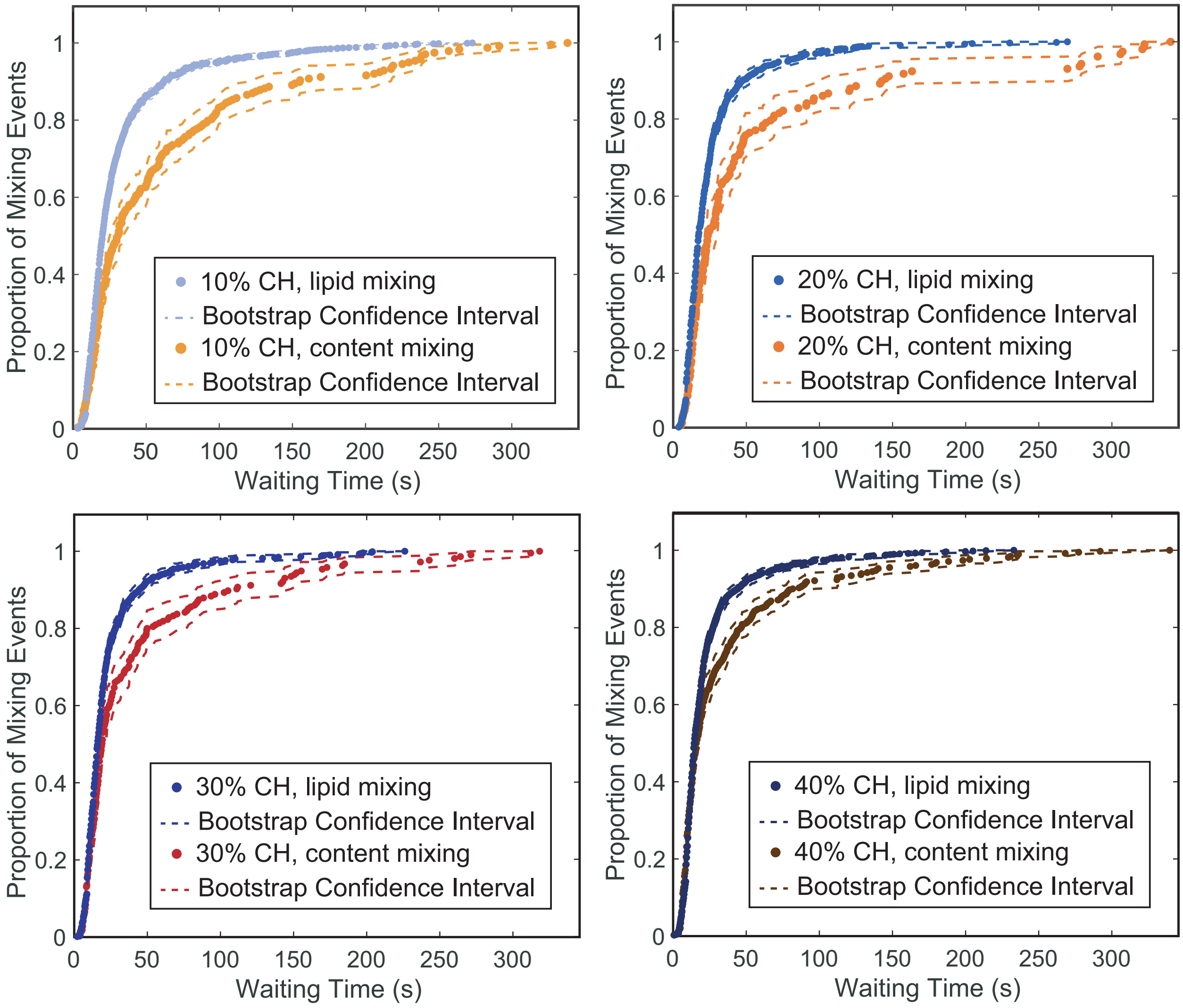


**Figure S4. Comparison between lipid and content mixing rates for various target membrane compositions**.

Target vesicles were tethered to IAV through GD1a. Content-labeled vesicles composed of 10% CH and all lipid-labeled vesicles were tethered through GD1a and DNA-lipid hybridization. Wait times from pH drop to lipid or content mixing for individual fusion events are plotted as CDFs. Lipid mixing traces were reproduced from reference (2) with permission. For all target vesicle compositions, content mixing is a slower process than lipid mixing (within bootstrap resampling error, 95% confidence intervals, dashed lines).

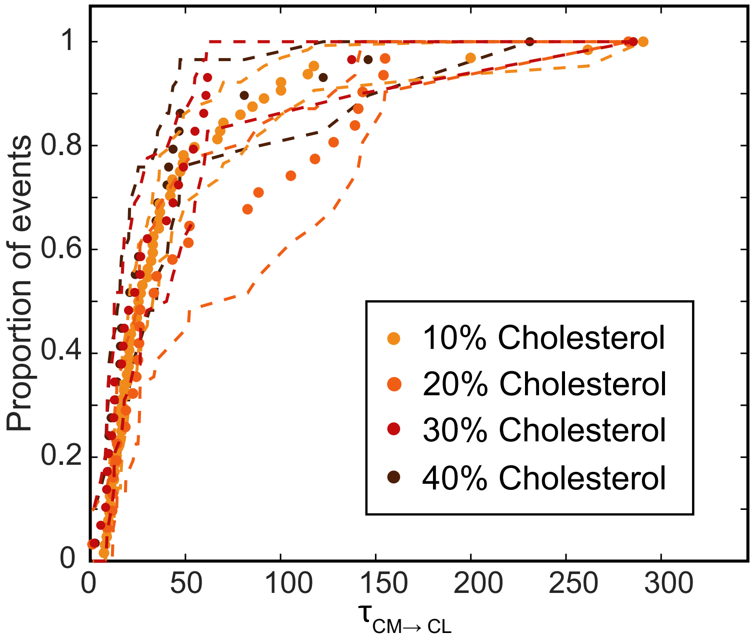


**Figure S5. Comparison of intervals from content mixing to content loss for vesicles containing various amounts of CH.**

$\tau_{CM\rightarrow CL}$ values for individual events are plotted as CDFs. $N_{CM\rightarrow CL}$ values for vesicles that contained a high fraction of CH were too low to observe any significant difference between compositions. Dashed lines represent bootstrap resampling error with 95% confidence intervals. $N_{CM\rightarrow CL}$ = 64 (10% CH), 31 (20% CH), 29 (30% CH), 29 (40% CH).

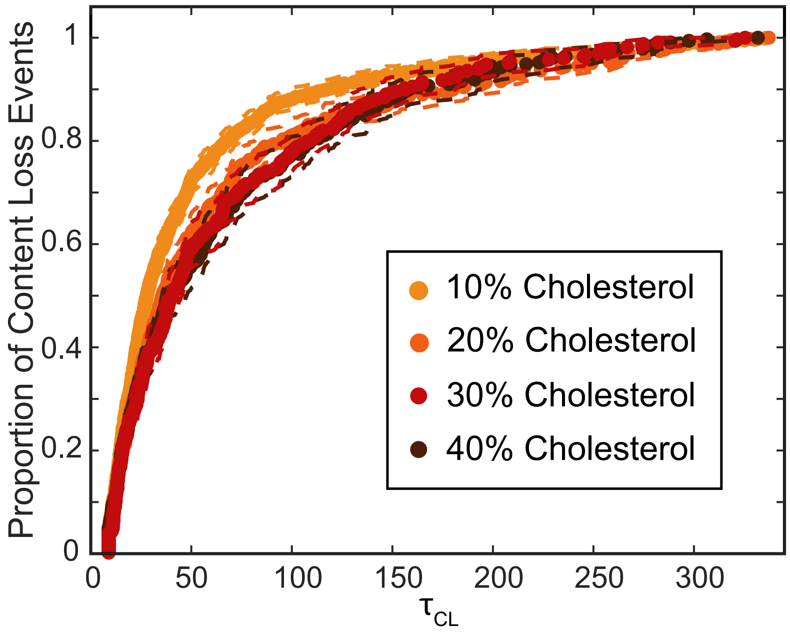


**Figure S6. The rate of content loss events is not affected by target membrane cholesterol.**
Wait times from pH drop to content loss ($\tau_{CL}$) for individual events are plotted as CDFs. For all target membrane compositions tested, the rate of content loss is the same within bootstrap resampling error (95% confidence intervals, dashed lines). As CH in target vesicles increases, there is no significant difference in the fraction of vesicles that undergo content loss: 724/5250 for 10% CH, 427/2458 for 20% CH, 328/2153 for 30% CH, and 333/2411 for 40% CH.

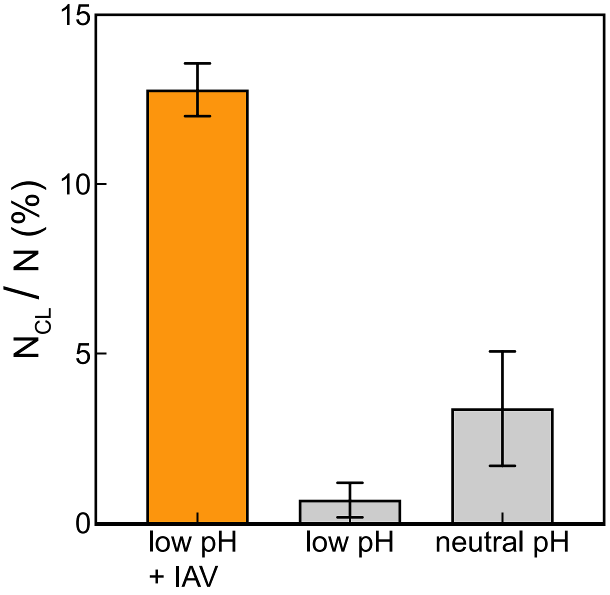


**Figure S7. There are significantly more content loss events at low pH when vesicles are tethered to IAV.**
In microfluidic flow cells, glass slides were functionalized as described in SI methods. For negative control experiments (gray), content-labeled vesicles (70% POPC, 20% DOPE, 10% CH) displayed DNA-lipids (sequence A') for surface tethering. Points represent the average frequency of content loss ± bootstrap resampling error**.** Due to the baseline of content loss events that take place when flow cells are exchanged with neutral pH buffer, all experiments were rinsed with a standardized amount of buffer.

*Single-virus binding assay to test for antibody coverage.* The design for this single-virus binding assay was based on previous studies (4, 5). 4 µL of TR-DHPE labeled IAV (2 mg/mL) were incubated with 1 µL of solutions of monoclonal antibodies (various concentrations, diluted in HB buffer) for 2 hours at room temperature (22˚C), then samples were kept on ice. In a flow cell, 1 µL of antibody-bound virions was introduced to a GD1a-displaying SLB and allowed to bind for 60 seconds. Unbound virions were rinsed away at 600 µL/min using vesicle buffer, and the resulting number of bound virions for each FOV was quantified through spot analysis of fluorescence micrographs using MATLAB scripts.

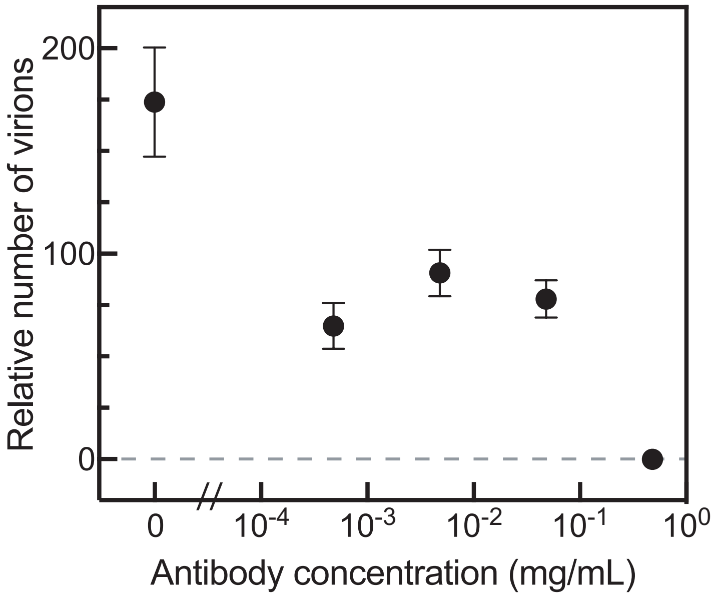


**Figure S8. Antibodies decrease IAV binding to a supported lipid bilayer displaying GD1a.** Points represent the average number of virions bound/FOV ± standard deviation. At least 35 images were taken for each concentration of antibodies.

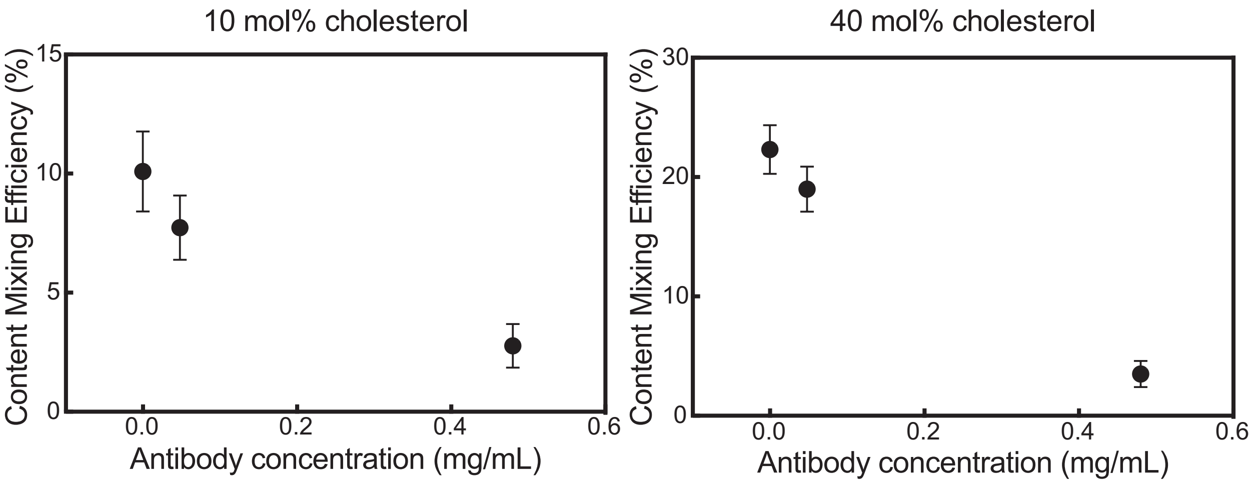


**Figure S9. Antibodies decrease the efficiency of content mixing to target vesicles containing 10 and 40 mol% cholesterol.**
Influenza A virions were incubated with various concentrations of monoclonal antibodies. Content-labeled vesicles composed of 10 and 40 mol% cholesterol were tethered to virions through DNA-lipid hybridization, and the pH was lowered to trigger content mixing and content loss. For both target membrane compositions, the addition of antibodies to IAV decreased the content mixing efficiency. Each data point represents at least 900 vesicles analyzed ± bootstrap resampling error.
